## Supplemental data for "Root cap cell corpse clearance limits microbial colonization in *Arabidopsis thaliana*"

### Supplemental Figures

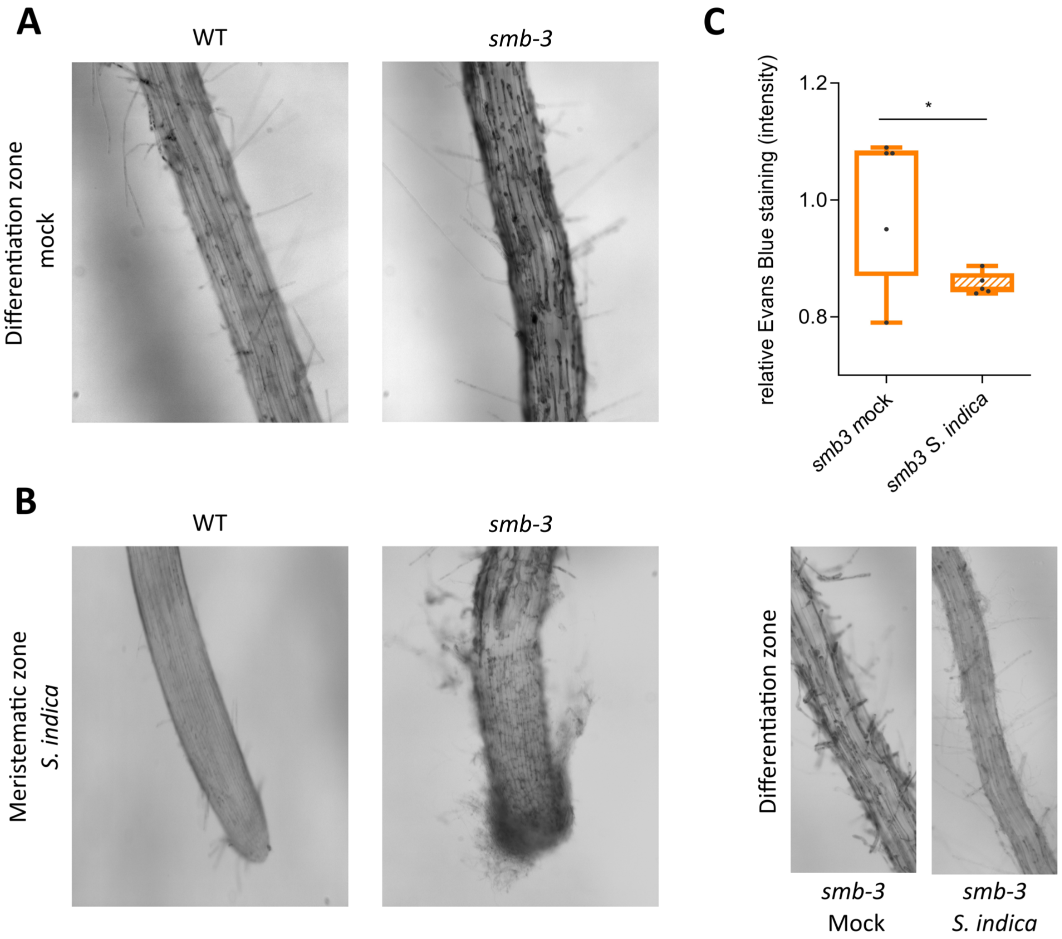


**Figure S1. Accumulation of LRC cell corpses on *smb-3* mutants.** (A) Evans blue staining of 14-day-old WT and *smb-3* mutant roots under mock conditions, showcasing the accumulation of LRC cell corpses along the differentiation zone caused by the SMB loss-of-function mutation. (B) Microscopy images of *S. indica* colonized WT and *smb-3* mutant roots visualizing the hypercolonization of the meristematic zone of *smb-3* mutant root tips at 10dpi, stained with Evans blue. (C) Quantification of cell death via Evans blue staining, showing a clearing of LRC cell corpses on *smb-3* mutant roots during *S. indica* colonization. 5 plants were imaged for each mock and *S. indica* treatment at 10 dpi. Statistical significance was determined using an unpaired, two-tailed Student's t-test before normalization ($F [4, 4] = 46.19$; $p < 0.05$). Microscopy images show corresponding mock and *S. indica*-treatments of *smb-3* mutant roots.

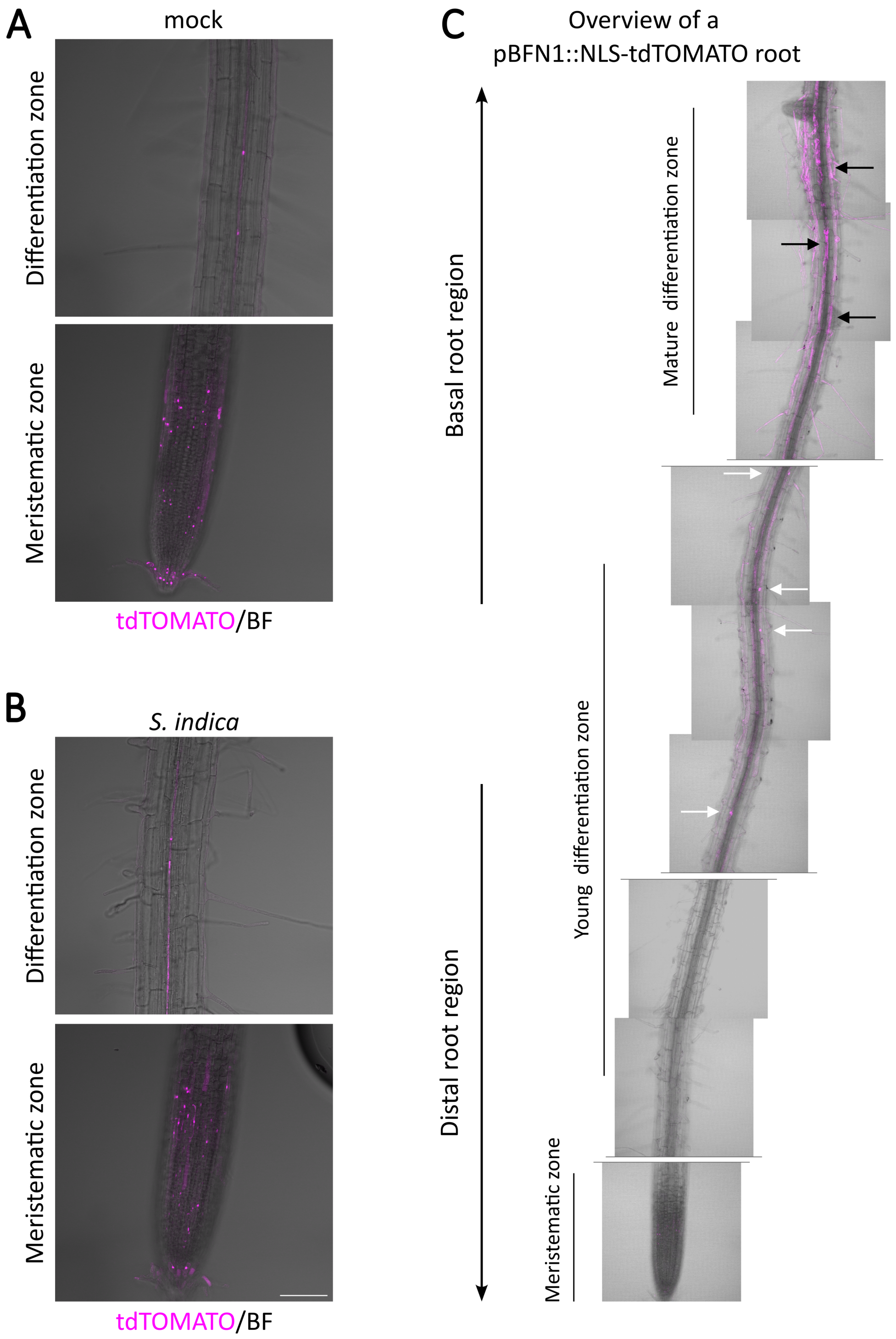


**Figure S2. Expression pattern of *BFN1* in root tissue**. CLSM images showing Z-stacks of the *BFN1* promoter reporter line pBFN1::NLS-tdTOMATO show *BFN1* expression via tdTOMATO signal (purple) accumulation in the nucleus of LRC cells and xylem cells. No significant changes in *BFN1* expression in the root cap and xylem were observed in mock treated (A) and *S. indica* colonized (B) roots at 7dpi. (C) Mosaic scan of a mock-treated pBFN1::NLS-tdTOMATO reporter plant. Overview shows single planes of the primary root axis. White arrows indicate *BFN1* expression in intact young epidermal cells, black arrows indicate *BFN1* expression in mature epidermal cells post nuclear rupture of a two-week-old seedling.

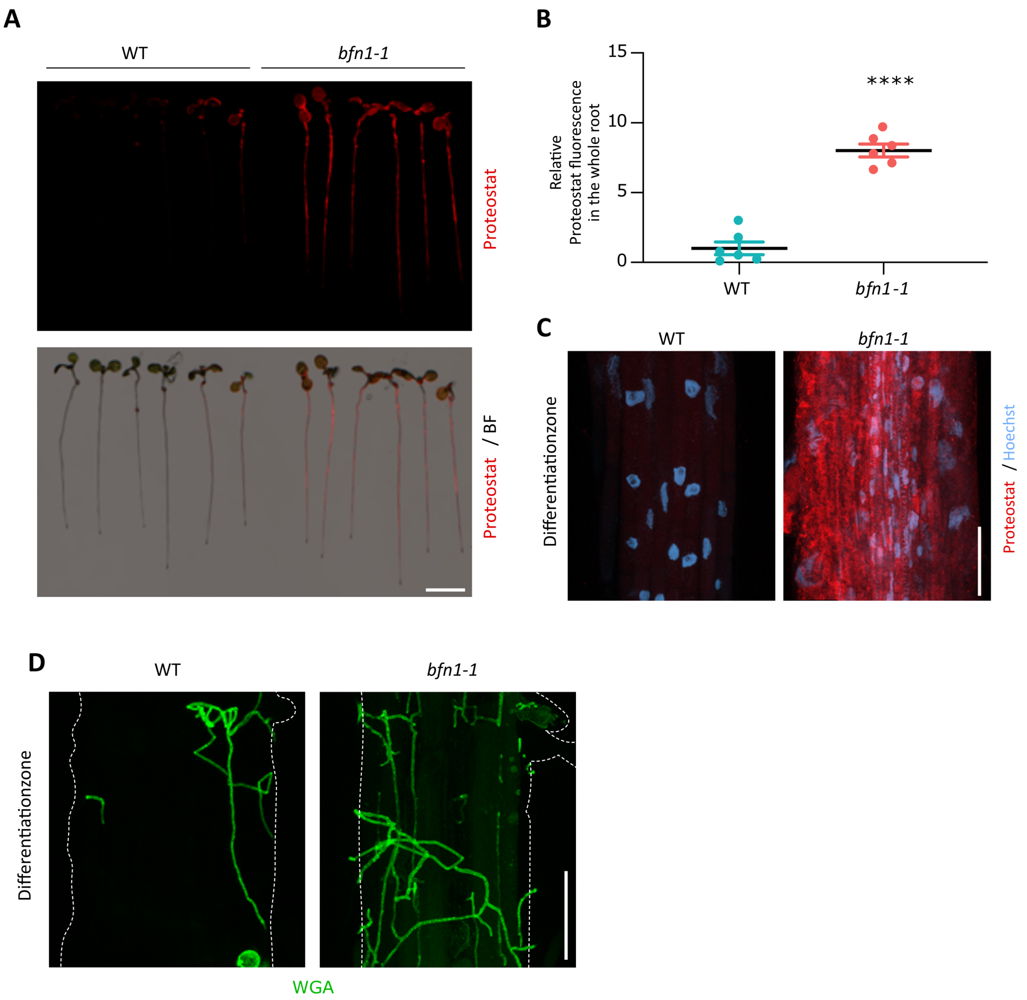


**Figure S3. *bfn1-1* mutant shows increased protein aggregation in roots.** (A) Images show Proteostat staining (red) of 10-day-old WT and *bfn1-1* seedlings. Roots were scanned and captured with a LI-COR Odyssey M imager using the bright field (BF) and the 520 nm wavelength channel. Scale bar indicates 5 mm. (B) Relative quantification of Proteostat signal in WT and *bfn1-1* roots. The statistical comparison was made by two-tailed Student's t-test for unpaired samples ($F [5, 5] = 1.005$; $p < 0.001$). (C) CLSM of differentiated cells of WT and *bfn1-1* roots. Proteostat (red) and Hoechst (blue) channels are shown. Scale indicates 50 µm. (D) CLSM images of the differentiation zone of Arabidopsis WT and *bfn1-1* roots inoculated with *S. indica* and stained with WGA-AF 488. Scale represents 50 µm.

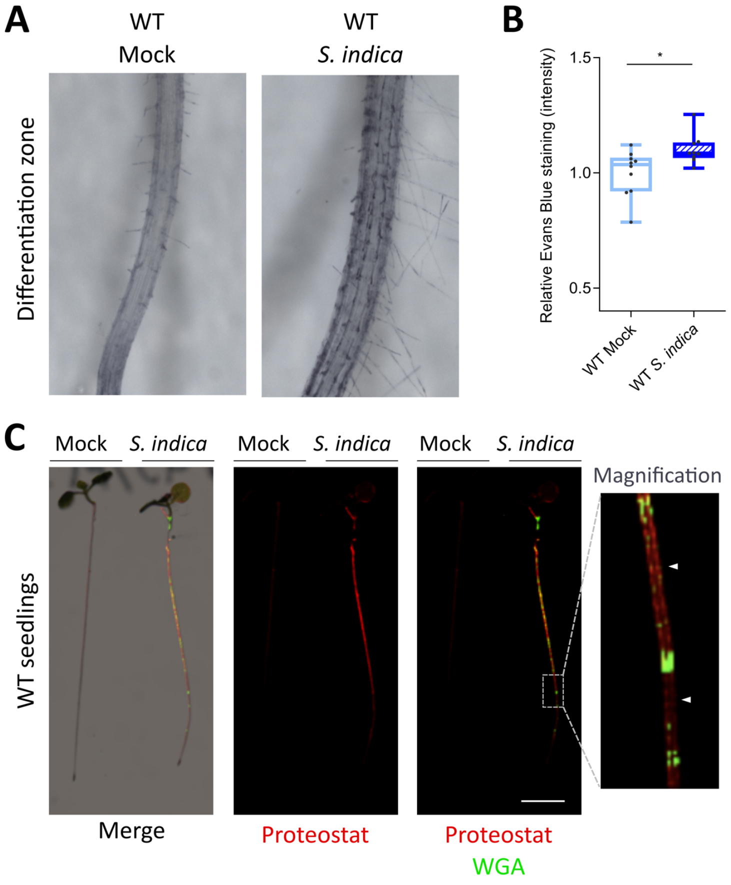


**Figure S4. Phenotypic analysis of *S. indica* colonization on WT Arabidopsis roots.** (A) Evans blue staining of WT roots visualize *S. indica*-induced cell death in the differentiation zone at 12 dpi. (B) Quantification of *S. indica* induced cell death in WT Arabidopsis, measuring relative Evans blue staining intensity (representative images shown in A). 10 plants at 12 dpi were evaluated. Statistical analysis was performed with a two-tailed Student's t test for unpaired samples ($F [9, 8] = 2.248$; $p < 0.05$). (C) Representative images of *S. indica*-colonized WT seedlings stained with Proteostat (red) and WGA-AF 488 (green), showing *S. indica*-induced protein aggregation/misfolding. Magnification panel shows Proteostat staining in zones of the root where *S. indica* is not present (white arrowheads). Scale indicates 5 mm.

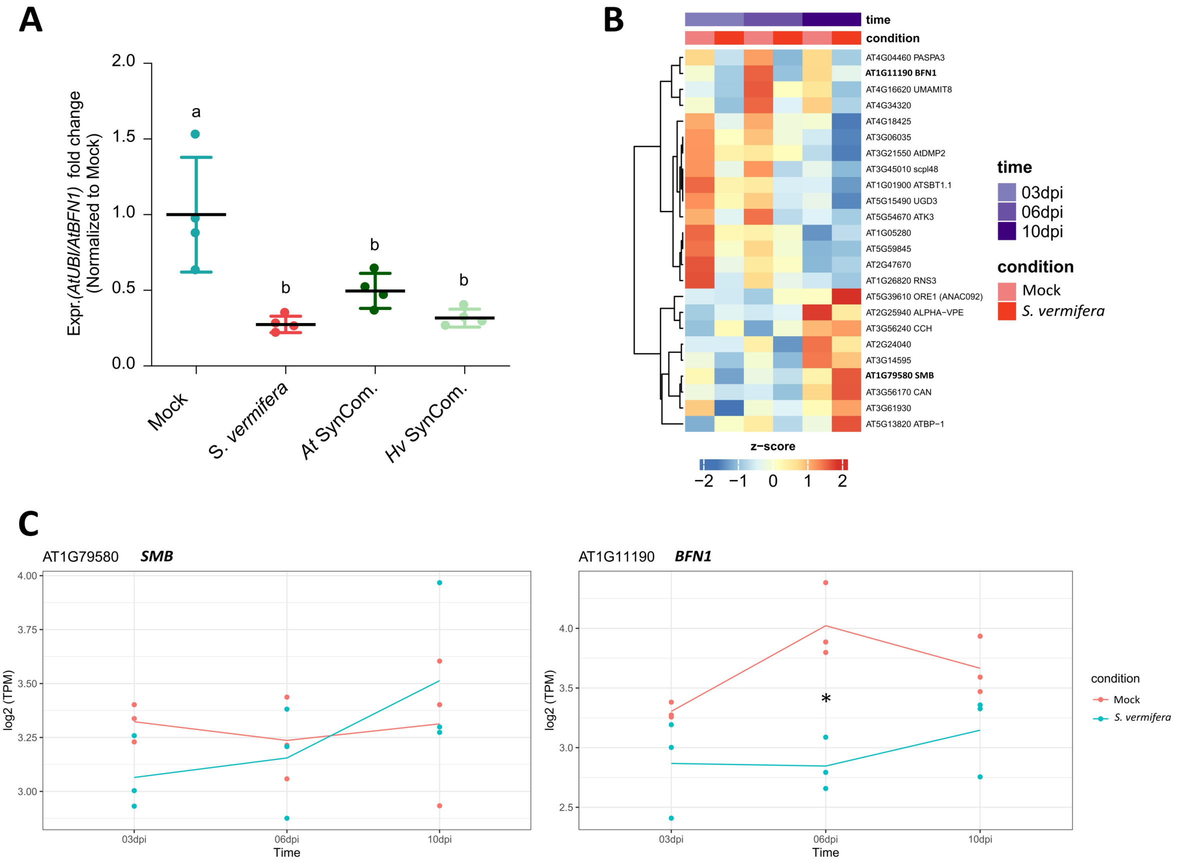


**Figure S5. *AtBFN1* is downregulated during colonization with beneficial microbes**. (A) qRT-PCR shows downregulation of *BFN1* in Arabidopsis during colonization by *S. vermifera* and bacterial SynComs from *H. vulgare* (Hv) and *A. thaliana*. RNA was harvested from 4 replicates at 6 dpi, pooling 30 plants per replicate for each treatment. *BFN1* expression levels are normalized to mock conditions. Statistical evaluation was performed via one-way ANOVA and Tukey's post hoc test ($F [3, 12] = 10.84$; $p < 0.001$). (B) Heatmap shows RNA-Seq expression of dPCD marker genes (Olvera-Carrillo et al., 2015) in mock-treated and *S. vermifera*-colonized Arabidopsis roots at 3, 6 and 10 dpi. TPM expression values are log2 transformed and row-scaled, showing an average of three biological replicates. Genes are clustered using Spearman correlation as distance measure. (C) *BFN1* and *SMB* expression profiles corresponding to RNA-Seq expression. Asterisk indicates significantly different expression (adjusted p-value < 0.05).

**Table S1. Primers used in this study**

| Primer ID | Sequence (5' to 3') | Target gene |
|---|---|---|
| AtUBI_F | CCAAGCCGAAGAAGATCAAG | AT3G62250 (*AtUBI5*) |
| AtUBI_R | ACTCCTTCCTCAAACGCTGA | AT3G62250 (*AtUBI5*) |
| FW_BFN1-A_qPCR | GGCGTCAAGTCTGGTGAAAC | AT1G11190 (*AtBFN1*) |
| RV_BFN1-A_qPCR | ACCCGGTTTAGTATCATGGCT | AT1G11190 (*AtBFN1*) |
| TEF_*S. indica*_qPCR_F | GCAAGTTCTCCGAGCTCATC | Transcription elongation factor |
| TEF_*S. indica*_qPCR_R | CCAAGTGGTGGGTACTCGTT | Transcription elongation factor |
